# Detecting cell type from single cell RNA sequencing based on deep bi-stochastic graph regularized matrix factorization

**DOI:** 10.1101/2022.05.16.492212

**Authors:** Wei Lan, Jianwei Chen, Qingfeng Chen, Jin Liu, Jianxin Wang, Yi-Ping Phoebe Chen

**Affiliations:** School of Computer, Electronic and Information, Guangxi University, Nanning, Guangxi, 530004 and Hunan Provincial Key Lab on Bioinformatics, School of Computer Science and Engineering, Central South University, Changsha, Hunan, 410083, China.; School of Computer, Electronic and Information, Guangxi University, Nanning, Guangxi, 530004, China.; School of Computer, Electronic and Information and State Key Laboratory for Conservation and Utilization of Subtropical Agro-bioresources, Guangxi University, Nanning, Guangxi, 530004, China.; Hunan Provincial Key Lab on Bioinformatics, School of Computer Science and Engineering, Central South University, Changsha, Hunan, 410083, China.; Department of Computer Science and Information Technology, La Trobe University, Melbourne Victoria 3086, Australia.

**Keywords:** ScRNA-seq clustering, Deep matrix factorization, Bi-stochastic graph regularization

## Abstract

The application of fruitful achievement of single-cell RNA-sequencing (scRNA-seq) technology has generated huge amount of gene transcriptome data. It has provided a whole new perspective to analyze the transcriptome at single-cell level. Cluster analysis of scRNA-seq is an efficient approach to reveal unknown heterogeneity and functional diversity of cell populations, which could further assist researchers to explore pathogenesis and biomarkers of diseases. In this paper, we propose a new cluster method (DSINMF) based on deep matrix factorization to detect cell type in the scRNA-seq data. In our method, the feature selection is used to reduce redundant features. Then, the imputation method is utilized to impute dropout events. Further, the dimension reduction is utilized to reduce the impact of noise. Finally, the deep matrix factorization with bi-stochastic graph regularization is employed to cluster scRNA-seq data. To evaluate the performance of DSINMF, eight datasets are used as test sets in the experiment. The experimental results show DSINMF outperformances than other state-of-the-art methods in clustering performance.

## 1 Introduction

SINGLE cell RNA sequencing (scRNA-seq) is an efficient approach to reveal unknown heterogeneity and functional diversity of cell populations, which could further assist researchers to explore pathogenesis of diseases. Cluster analysis of scRNA-seq is a critical way to discover the heterogeneity of cells. Detecting cell types from single cell RNA sequencing data is an unsupervised learning task. There are some challenges for this task. Firstly, the noise may make the performance of many existing approaches less than expected [1]. Second, the high dimension of scRNA-seq data also leads to poor performance on many clustering algorithms based on similarity. In addition, sparsity is a significant characteristics of single cell data, in other word, scRNA-seq data have a large number of zero entries. It also restricted the application of cluster method in single-cell data analysis [2], [3].

In recent years, several approaches have been proposed for clustering single cell transcriptome data. Chen et al. [4] introduced a computational method (SNN-Cliq) to cluster scRNA-seq data based on shared nearest neighbor. It obtained the affinity matrix based nearest neighbor and cluster it by merging quasi-cliques. Satija et al. [5] presented a computational method (Seurat) to detect cell type based on the Shared Nearest Neighbors (SNN) graph. Wang et al. [6] proposed a cluster method (SIMLR) for scRNA-seq analysis based on multiple kernel learning. Park et al. [7] presented a method (MPSSC) to cluster scRNA data by utilizing multiple doubly stochastic affinity matrices to construct a robust similarity matrix. Kiselev et al. [8] designed a method (SC3) for scRNA-seq analysis. It transformed the original matrix into a number of matrices with different dimensions through spectral clustering, and then used principal component analysis to obtain cluster results. Zheng et al. [9] presented a method (SinNLRR) based on subspace clustering for scRNA-seq analysis. It measures the similarity between samples by using linear representation. Wu et al. [10] developed a method (DRjCC) based on matrix factorization to detect cell type.

Although these methods have achieved great successes in scRNA-seq cluster. There are still a great number of limitations. Firstly, the high dimensional data makes the ensemble learning method inefficient as computational consumption. Secondly, with the increasing of scRNA-seq data, some methods are unable to effectively extract the deep features of the data. Finally, some methods ignore the dropout problem of scRNA-seq, which may bring some bias of clustering result.

In this paper, we proposed a new method based on deep matrix factorization (DSINMF) for single cell cluster analysis. We summarize the main contributions of this paper as follows:

a. The feature selection can remove redundant features (genes), and select highly discriminant features (genes) for clustering according to the importance of features (genes).
b. We use imputation of matrix to overcome the limitation of dropout of the original data due to measurement errors which can promote the accuracy of clustering result.
c. The structure of multi-layer matrix factorization is utilized to extract the deep hidden features of biological data which can obtain the features in different layers.
d. The adaptive graph learning is used to learn high quality graph by using the bi-stochastic matrix. Further, the bi-stochastic graph regularization deep matrix factorization is proposed to obtain the result of cluster.
e. The partial parameters in DSINMF are updated with adaptive manner. Therefore, the user need not to set parameters manually.

## 2 METHODS

The flowchart of the DSINMF is shown in Figure 1. First, the feature selection method is used to reduce redundant features. Then, the dropout imputation is conducted to overcome the limitation of dropout. Further, the dimension reduction is employed to reduce the effect of noise. Finally, the bi-stochastic graph regularization deep matrix factorization is utilized to cluster the scRNA data by capturing the features of different subspace.

**Fig. 1.**
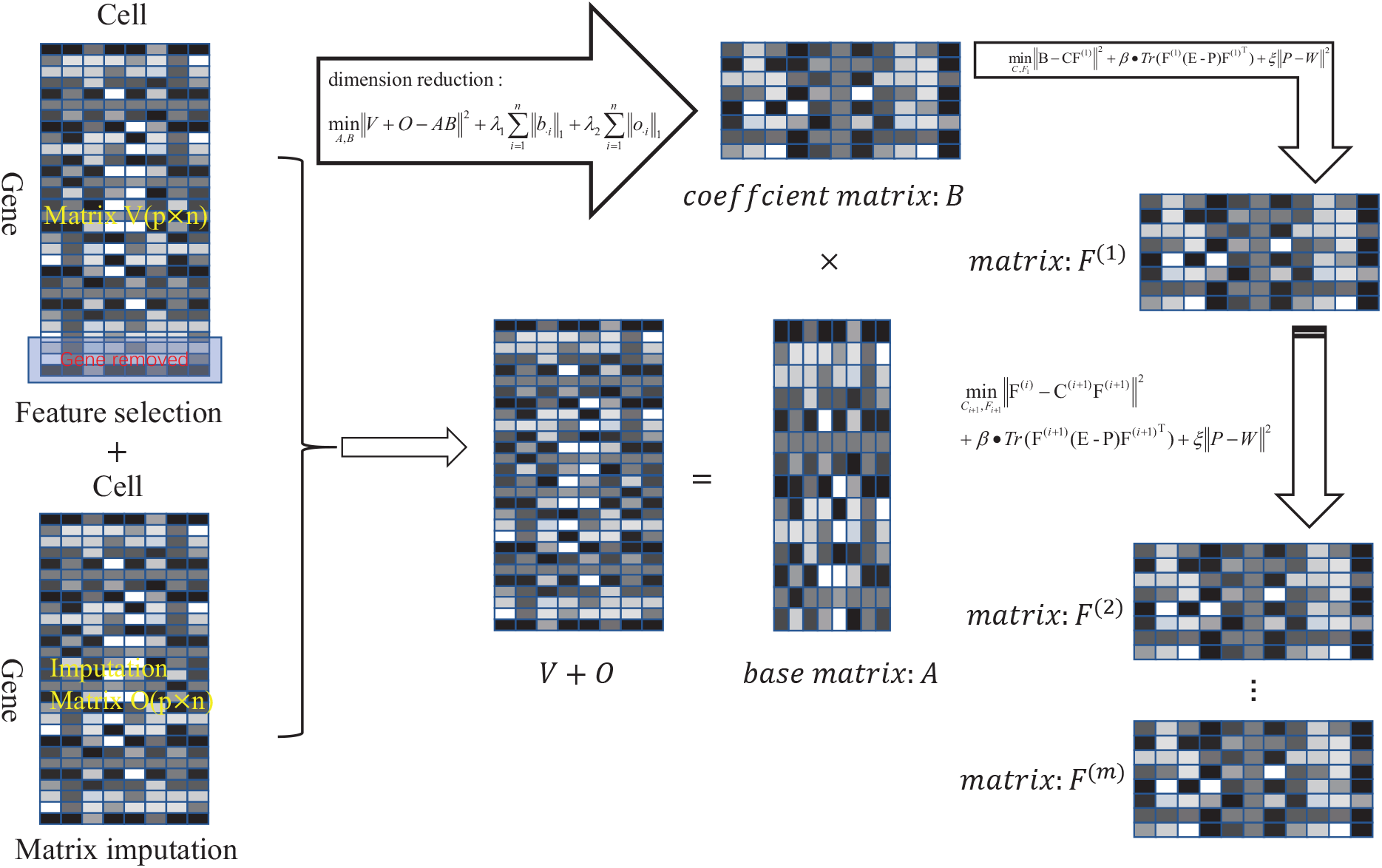
The flowchart of the DSINMF. First, some genes are removed by genetic screening, then matrix factorization is used to reduce the dimension of the data, and finally deep matrix factorization is used to extract the deep features of the data. where V represents the original data matrix and O represents the imputation matrix.

### 2.1 Feature selection

Single cell RNA-seq data clustering usually calculates the similarity between cells in graph regularization constraints. The feature selection can affect the quality of the similarity matrix, which consequently effects the performance of clustering. CaFew [11] is used to obtain the weight matrix 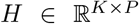. for *P* features (genes) in *K* clusters. The *CV* (coefficient of variation, defined as the ratio of the standard deviation to the mean) is used to measure the variation of feature weights across clusters, and further to select the “marker” genes for clustering. The linear model is established: *log*(*CV*^2^) = *a* ∗ *log*10(*mean*(*H_p._*)) + *b* to fit the CV by mean. After the fitting is completed, each gene will get a residual *d_p_*, which represents the difference between the true CV and the fitted value. These residuals also represent the difference between the weight coefficient change law of gene. Then, the residual value is normalized as: 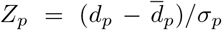, where *σ_p_* denotes the standard deviation of *H_p_*. *Z_p_* can reflect the importance of the gene in the cluster process, The smaller value of *Z_p_* is, the important of the gene. Therefore, the *Z_p_* is used as the criterion for feature selection.

### 2.2 Dimension reduction and dropout imputation

The non-negative matrix factorization (NMF) decomposes the given data matrix into two non-negative matrices. The original matrix factorization can be defined as follow:

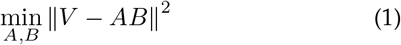

where ∥∗∥ denotes the Frobenious norm. the matrix *V* ∈ *R^P×n^* is constructed to represent the gene-by-cell expression data, where p and n denotes the number of genes and cells, respectively. *A* ∈ *R^p×r^* and *B* ∈ *R^r×n^* denote two non-negative sub-matrices after decomposition. r denotes the dimension of the subspace.

Due to the high dimension of raw data, the similarity of cells generated by using spatial distance or correlation measurements is not useful in distinguishing the categories of samples. Therefore, we use NMF to acquire low-dimensional representation for given data matrix *V*. In single cell transcriptomic data, sparse representation can improve the inter-pretability and accuracy of the algorithm [12]. Thus, the *L*_1_-norm is used to ensure the sparsity of data. The objective function is defined as follows:

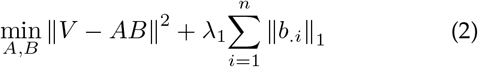

where *b_.i_* represents the column vector of *B*, ∥∗∥_1_ represents the *L*_1_-norm. *λ*_1_ denotes the penalty parameter for sparse control. In addition, there are many zero entries in the genecell expression matrix, which would be appeared when the dropout occurs [3]. To overcome this limitation, the matrix *O* ∈ *R^p×n^* is utilized to impute the dropouts where *O_ij_* = 0. Then, the original data matrix can be rewritten as the dropout recovery matrix *V* + *O*, which can be decomposed approximately into two sub-matrices *A* and *B* (*V* + *O* ≈ *AB*). It is reasonable to assume that *O* is sparse as a lot of zero entries in matrix. Therefore, the object function can be defined as follows:

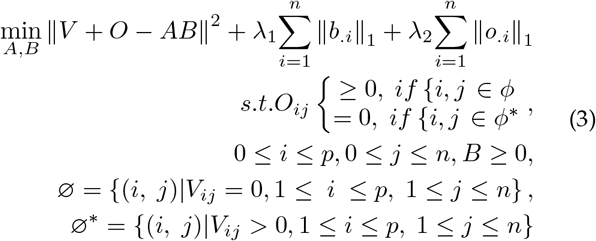

where *λ*_1_ and *λ*_2_ are the penalty parameters for sparse control.

### 2.3 Deep matrix factorization

In NMF, the matrix V is decomposed to two sub-matrices A and B, where the matrix A represents the feature matrix of the data sample. The matrix B represents the combination coefficient matrix that combines the features into the original data, and it also regarded as low-dimensional mapping data. In order to obtain the hidden features, the deep matrix factorization is used to extract the deep features of biological data. Finally, we extract all the features into the feature representation matrix 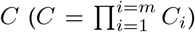, and the resulting matrix *F* is the coefficient of the combined features [13]. It is defined as follows:

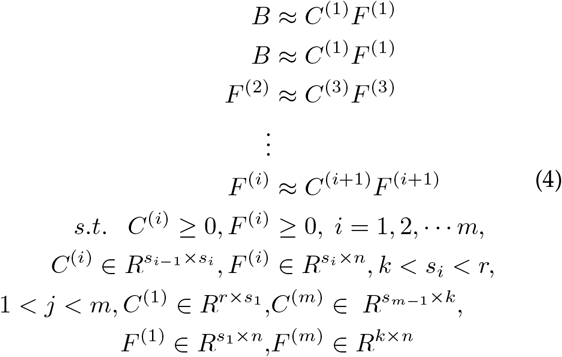

where *k* denotes the number of clusters. *m* denotes the total layer of decomposition. For i-th layer, the resulting matrix F(i) is input to the next level of decomposition. And finally, the resulting matrix *F*^(*m*)^ is obtained. Considering that the NMF is unable to detect the geometric space manifold structure contained in the high-dimensional data. Therefore, the projecting representation of the potential low-dimensional space may destroy the geometric structure of the original space [13], [14]. If the sample have high similarity in the original space, there is still a high correlation in the potential low dimensional projection space. Therefore, the inheritance of geometric structure should be obtained in the decomposition of each layer. The objective function of single-layer decomposition is defined as follows:

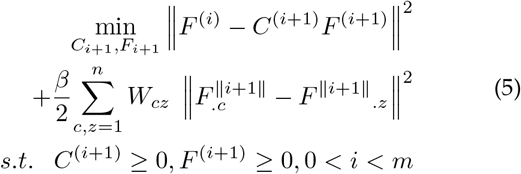

where m denotes the total layer of decomposition. *β* is the penalty coefficient for the regularization term. *C_.i_* and *F_.i_* are the vectors from the i-th column of *C* and *F*, respectively. *C*^(*i*)^ *and F*^(*i*)^ are the matrices obtained from the *i*-th decomposition. The affinity matrix *W* ∈ *R^n×n^* is calculated by using Cosine correlation coefficient, where each element denotes the similarity of pairwise cells. In addition, Cai et al. [15] have proved that the inheritance of a local topology structure can be represented as trace optimization, i.e.

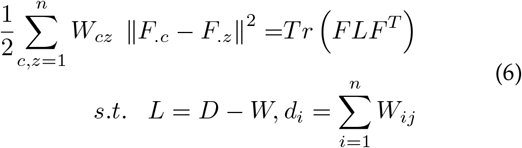

where *L* is the Laplacian matrix of graph *W*. *D* is a diag matrix. Thus, the Eq. (5) can be defined as follow:

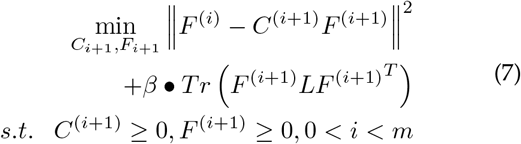

### 2.4 Bi-stochastic graph regularized matrix factorization

It is not appropriate to directly use the similarity matrix to ensure the manifold structure constraint on the matrix factorization. The graph constructed by K-NN (K-Nearest Neighbor) cannot represent the original data structure. Therefore, the bi-stochastics graph regularized [16] is used to learn an adaptive graph *P*. The *P* should have two properties as follows: firstly, *P* can describe the probability of any two samples in the same cluster. Therefore, the sum of each of its rows is 1, and each value in the matrix must be non-negative. Secondly, *P* must satisfy the property of symmetry as it represents the similarity of each pair of cells. In addition, the constraint *diag*(*P*) = 0 is added as each vertex should not be connected by itself. In the end, the bistochastic matrix is obtained from the input graph *W* by solving the following problem:

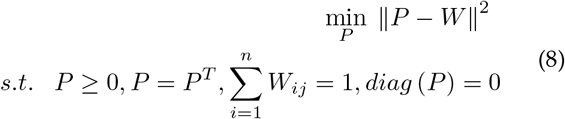

Combining the Eq. (7) and Eq. (8), the bi-stochastic graph regularized matrix factorization is defined as follow:

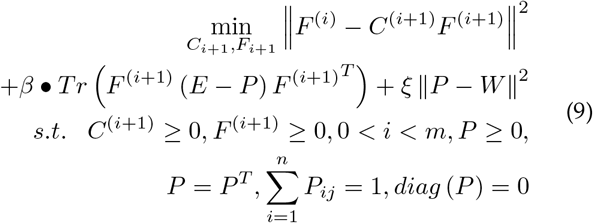

### 2.5 Optimization of DSINMF

The objective function of this model is a non-convex function, naturally, the alternating direction method of multipliers (ADMM) [17] can be applied to solve optimization problem.

#### Update P

In order to solve Eq. (8), it is divided into the following two sub problems:

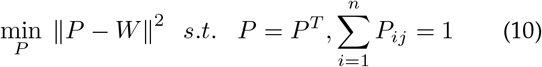

and

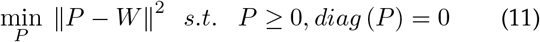

Eq. (10) can be solved as follow:

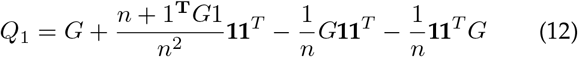

where 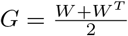, **1** is a vector with all elements 1. n is the size of *W*.

Eq. (11) can be solved as follow:

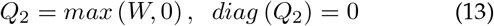

We view *Q*_1_ as *W* of Eq. (13) and *Q*_2_ as *W* of Eq. (12), thus *P* can be obtained by alternatingly updating the variable sets *Q*_1_ and *Q*_2_.

#### Update A

Let Θ ∈ *R*^*s*_1_×*n*^ be the Lagrange multiplier for *B* ≥ 0, and then the Lagrange *L* of DSINMF objective function is defined as follow:

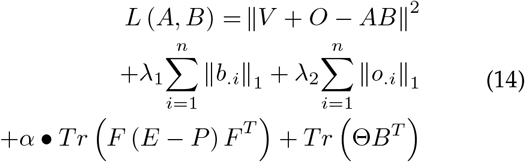

Firstly, the partial derivatives of *L* with respect to *A*

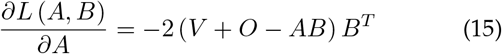

let 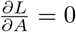, we can get the update of *A* is

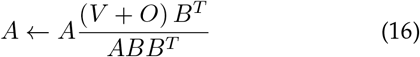

#### Update B

For variable *B*, we can get loss function of it as

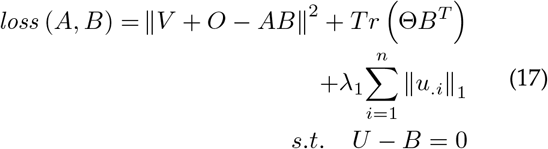

where *u_.i_* is the i-th column of matrix *V*. We use ADMM to optimize Eq. (17), it can be redefined as follow:

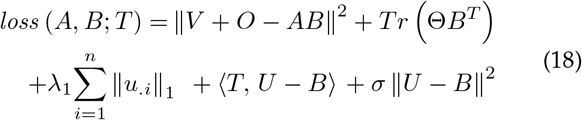

where *σ* > 0 is the penalty parameter, *T* ∈ *R^r×n^* is Lagrange multiplier. Then, the partial derivative of *loss* (*A, B*; *T*) with respect to B:

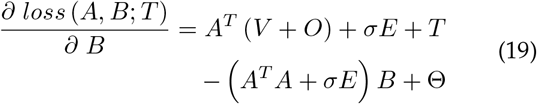

using the KKT condition Θ*B^T^* = 0. Let 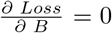, we can get the update of *B* is

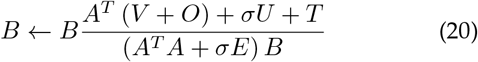

where *E* is identity matrix.

#### Update parameter T

For variable *T*, we can get update

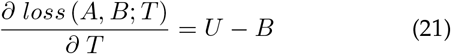

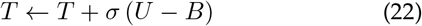

#### Update parameter U

*U* can be approximately by the Eq. (18), then

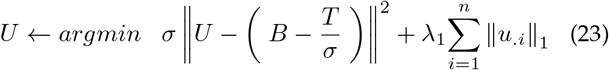

#### Update parameter O

*O* can be obtained by the Eq. (14), then

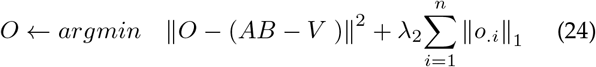

Both Eq. (23) and Eq. (24) are soft threshold problems, so the update mode of *A* and *B* is

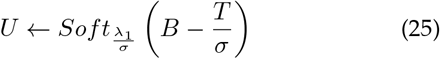

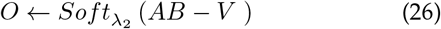

where *Soft_τ_*(*x*) is a soft threshold function, it can be defined as

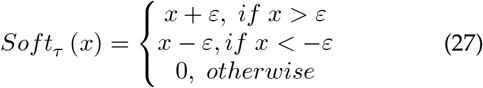

#### Update matrix C

For variable *C*, we can get

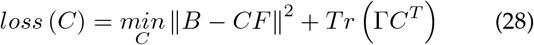

then, the partial derivatives of function with respect to *C*

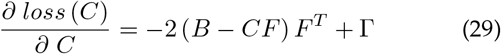

using the KKT condition Γ*C^T^* = 0. Let 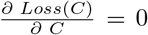, we get the update of *C* as follow:

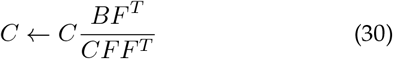

#### Update matrix F

For variable *F*, we can get loss function of it as

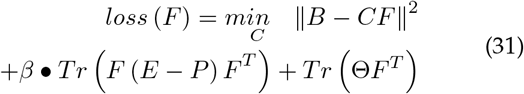

Then, the partial derivatives of function with respect to *F*

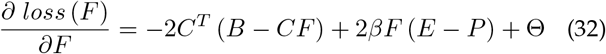

using the KKT condition Θ*F^T^* = 0. Let 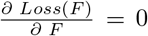, we get the update of *F* as follow:

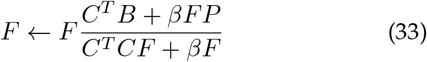

According to Eq. (30) and Eq. (33), we can get the update of Eq. (9) as follows:

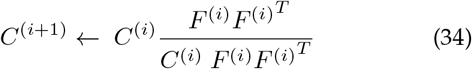

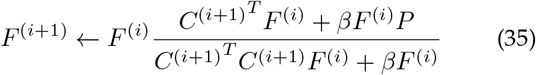

where *F*_1_ = B. Based on the above derivation, the algorithm is shown in Algorithm 1 as follow:

##### Algorithm 1: The algorithm of DSINMF

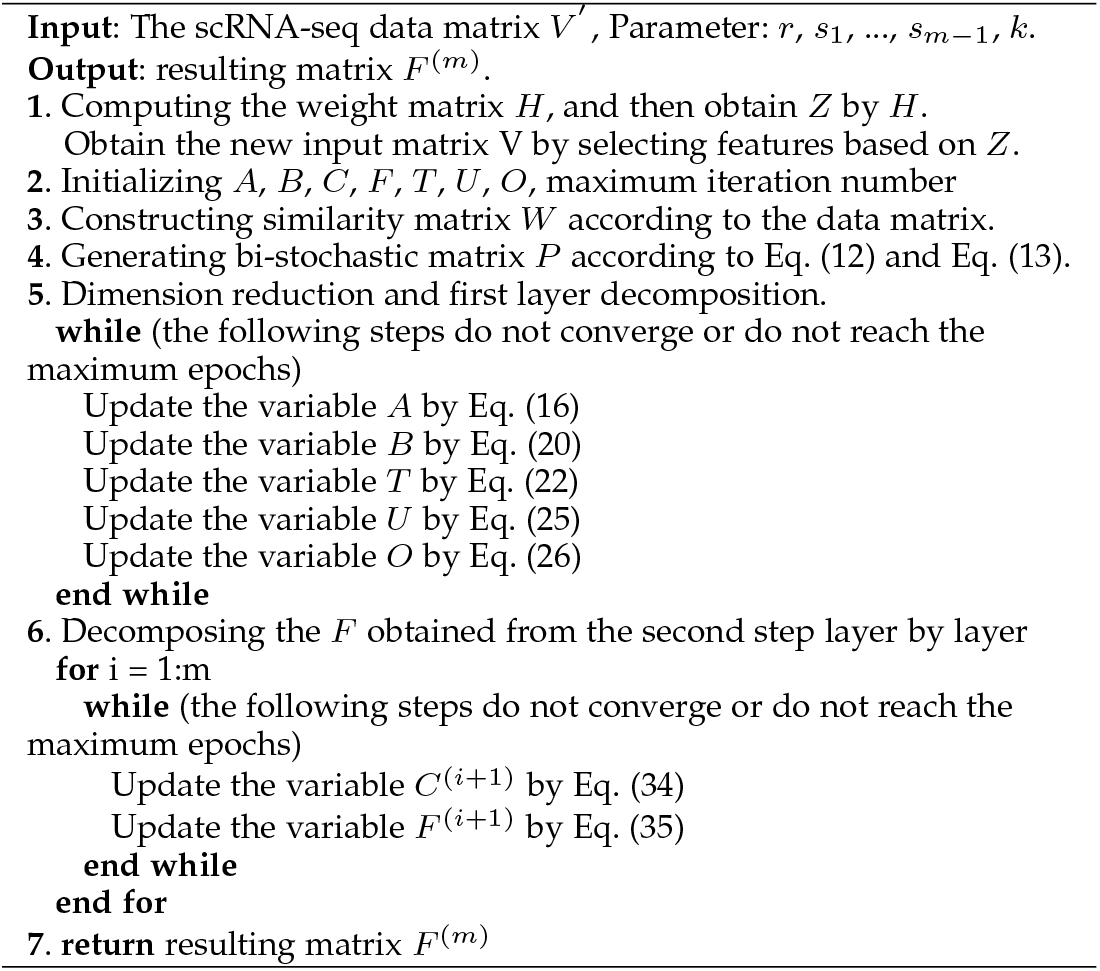

### 2.6 Datasets and Evaluation Metrics

Eight datasets of human and mouse single-cell RNA sequencing have been collected for our experiment including five datasets from human (Pollen [18], Chu [19], Patel [20], Zheng [21], Darmanis [18]) and three datasets from mouse(Goolam [22], Haber [23] and Mouse2 [21]). All original data are processed by using the log transformation conversion. The detailed descriptions of the datasets are shown in Table 1.

**TABLE 1.** The description of datasets used in experiments.

| Dataset | Cells | Genes | Types | Species | Ref |
| --- | --- | --- | --- | --- | --- |
| Pollen | 249 | 10015 | 11 | Homo sapiens | Pollen et al., 2014 |
| Chu | 1018 | 19097 | 7 | Homo sapiens | Chu et al., 2016 |
| Patel | 430 | 5769 | 5 | Homo sapiens | Patel et al., 2014 |
| Zheng | 500 | 4776 | 3 | Homo sapiens | Zheng et al., 2017 |
| Goolam | 124 | 40315 | 5 | Mus musculus | Goolam et al., 2016 |
| Haber | 1522 | 20108 | 9 | Mus musculus | Haber et al., 2017 |
| Darmanis | 420 | 22085 | 8 | Homo sapiens | Darmanis et al., 2015 |
| Mouse2 | 1064 | 14878 | 13 | Mus musculus | Biase et al., 2014 |

Three evaluation metrics including Adjust Rand Index (ARI) [24], Normalized Mutual Information (NMI) [24] and Adjust Mutual Information (AMI) [25] are utilized to measure the performance of cluster method. The ARI is widely used to evaluate the effect of single cell type detection. The cluster division *K* = (*k*_1_, *k*_2_, *k*_3_,…, *k_i_*) of the whole samples is obtained through cluster method, and then the data set provides the standard division *C* = (*c*_1_, *c*_2_, *c*_3_,…, *c_j_*) ofall samples, ARI is defined as follows:

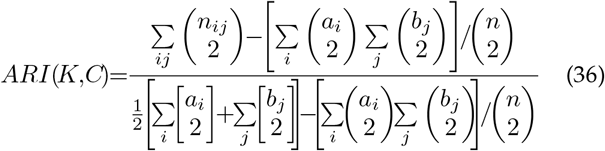

where n is the size of sample set, 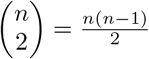, *n_ij_* denotes the number of cells shared by cluster *k_i_* and cluster *c_j_*, *a_i_* is the size of *k_i_*, and *b_j_* is the size of *c_j_*.

The NMI and AMI are also utilized to measure the distribution similarity between the two partitioning of clustering. The NMI and AMI are defined as follows:

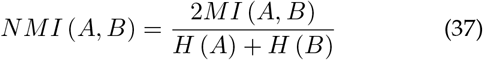

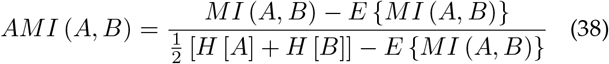

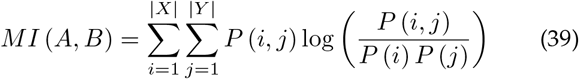

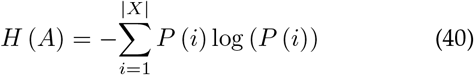

where |*X*| = *size*(*unique*(*A*)), |*Y*| = *size*(*unique*(*B*)), *i* ∈ *unique*(*A*), *j* ∈ *unique*(*B*), *P* (*i*) and *P*(*j*) represent the probabilities of *i* and *j* appearing in *A* and *B*, respectively. *P*(*i, j*) is the probability that *i* and *j* are both in the same cluster.

## 3 RESULTS

We compare our model with six other state-of-art clustering methods which are shown as follows:

**SNN-clique [4]**: SNN-clique is a method based on shared neighbor to measure the similarity between cells. This method aims to construct a more stable matrix of intercell relationship.
**K-means [26]**: the most commonly used clustering method.
**MPSSC [7]**: MPSSC adopts similar matrix fusion method, in which the key is to calculate the multi-kernels similarity between the cells.
**NMF [27]**: it is the conventional non-negative matrix factorization.
**pcaReduce [28]**: it firstly conducts dimensionality reduction processing (PCA) on the original data matrix, then adopts the idea of bottom-up hierarchical clustering, and replaces the process of merge one by one with the autoencoder.
**Seurat [5]**: a computational method based graph clustering to infer cellular localization by integrating single-cell RNA-seq data with RNA patterns in situ.
**DRjCC [10]**: it is an improved method based on NMF by using two-tier matrix factorization.

### 3.1 Comparative analysis of clustering performance

In the experiment, eight scRNA-seq datasets are selected as gold standard datasets, which are described in Table 1. All datasets are derived from human and mouse. For fairness, we provide the true number of clusters to K-means [26], MPSSC [7], pcaReduce [28], Seurat [5] and DRjCC [10] while SNN-Cliq [4] cannot be set to a certain number of clusters, and other parameters are set to default. We use K-means [26] with Pearson similarity as a baseline method. Fig. 2 summarizes the NMI, AMI and ARI of these methods on the eight datasets. The DSINMF achieves the best performance in six datasets, and the performances on the other two datasets are comparable with the best method. It can be found from Fig. 2A that compared with other algorithms, the DSINMF achieves the best performance on six datasets (Haber, Patel, Goolam, Chu, Patel and Mouse2). Specially, the DSINMF obtains 35% improvement (compare to DRjCC base on NMI) on dataset Haber (NMI: 0.588, DRjCC: 0.433), the improvements on other data sets are: Patel (21%), Mouse2 (11.2%), Darmanis (1%), Goolam (7%) and Chu (2%). Fig. 2B and 2C show the clustering performance in term of AMI and ARI respectively. It can be discovered that DSINMF also achieves the great performance on all datasets. In addition, Table 2 shows the detail numerical results of each algorithm on eight datasets. The fig. 3 shows the twodimensional visualization results of DSINMF and DRjCC by using t-SNE on four datasets (Haber, Patel, Mouse2, Goolam). It can be observed that the cluster distribution obtained by DSINMF is more significant than that obtained by DRjCC. For example, the dividing lines between different clusters of DSINMF are more obvious than DRjCC in Haber.

**Fig. 2.**
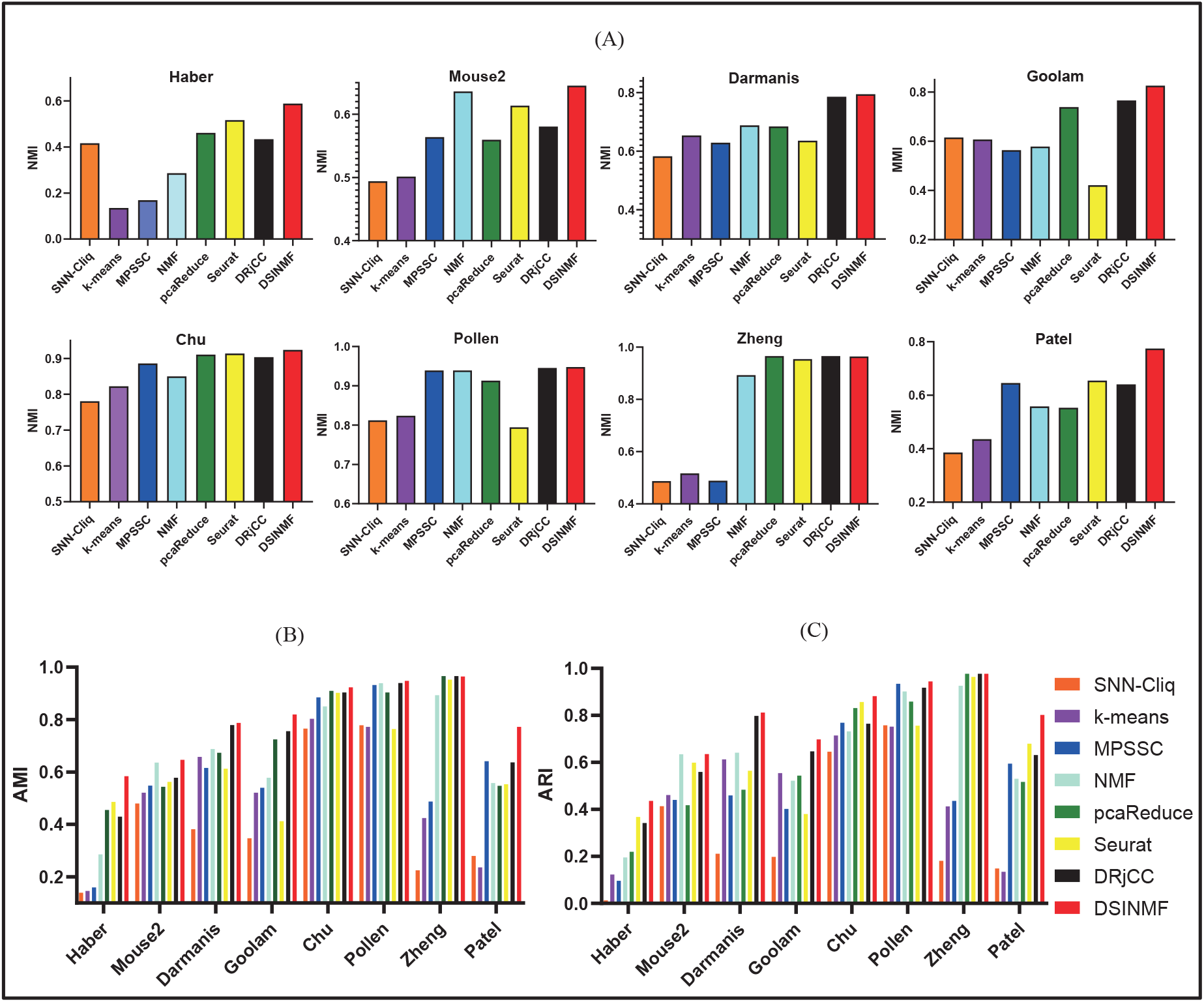
The performance of different algorithms on eight scRNA-seq datasets in term of three measurements: (A)NMI, (B)AMI and (C)ARI. The numerical results of the experiment are the average of ten operations.

**Fig. 3.**
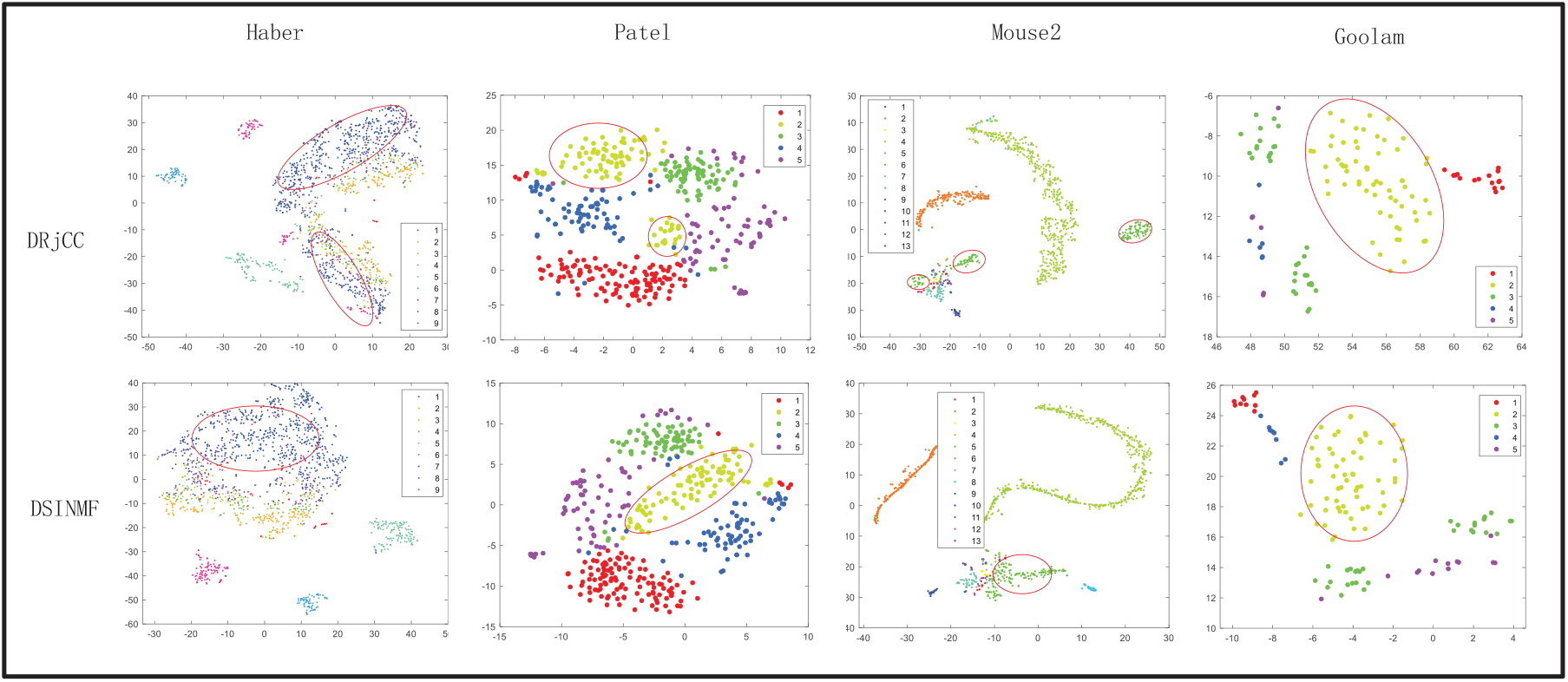
The visualization of origin data, DSINMF and DRjCC by using t-SNE.

**TABLE 2.**
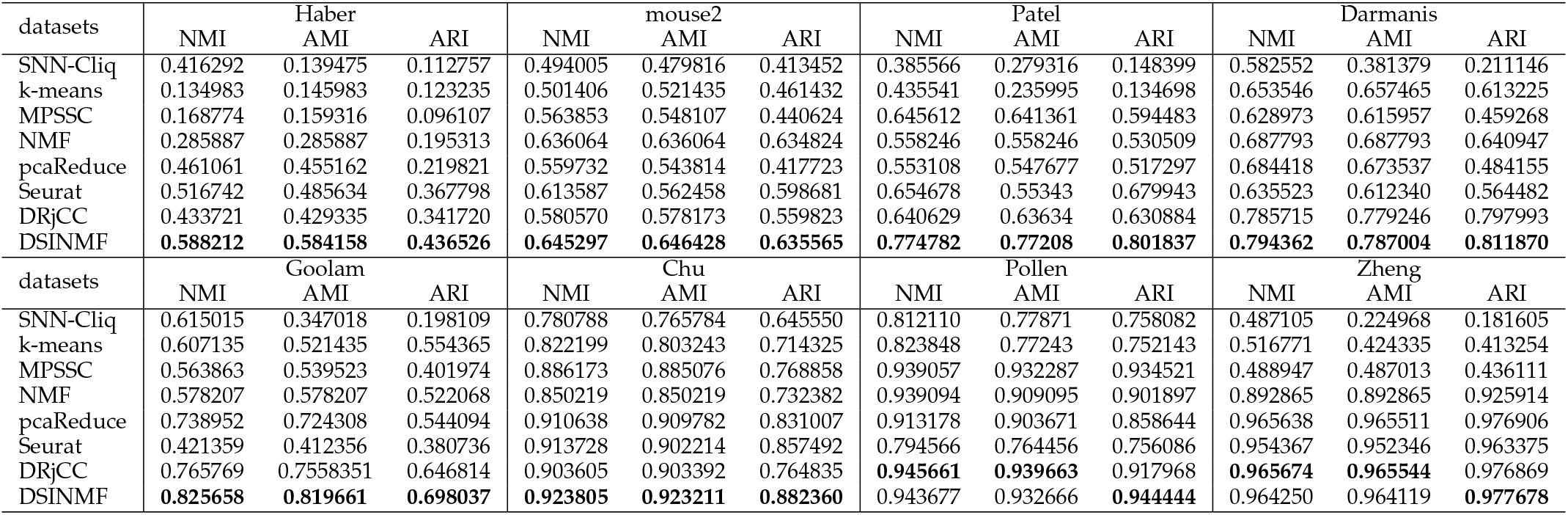
The detail value (NMI, AMI and ARI) of different methods.

### 3.2 Parameter selection

In DSINMF, the parameter *λ*_1_ and *λ*_2_ are the penalty factors of matrices *O* and *B*, respectively. In addition, the parameter *β* is the penalty factors of graph regularization. Similar to Shah [27], the parameters *λ*_1_ and *λ*_2_ and *β* are updated with adaptive manner during iteration. It is defined as follows:

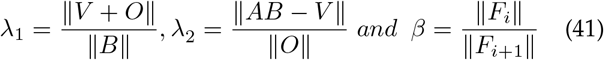

In feature selection part, the *Z_p_* is used to rank the features (genes). The top percentage method is used for feature selection based on the rank of features (gene). The result is shown in Fig. 4A. It can be found that the cluster performance is the best when the selected feature between 50% an 70%.

**Fig. 4.**
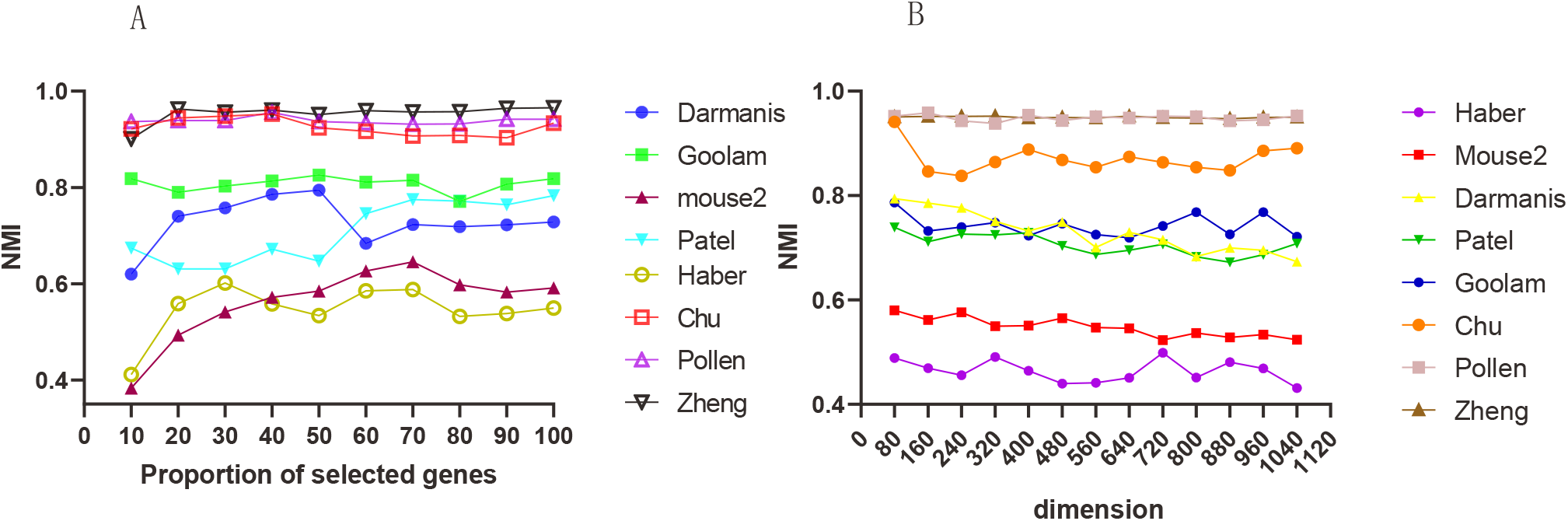
(A) The effect of proportion of selected genes in feature selection. The value on the horizontal axis represents the percentage of selected genes in the set of candidate genes, and the selected genes are ranked higher by default. (B) The effect of different dimension in dimension reduction. The value on the horizontal axis represents the target dimension of the dimensionality reduction layer. Here, the effect change of the eight datasets under the change of the target dimension is shown.

To test the effect of dimension in dimension reduction, we set the dimension of first layer arrange from 80 to1040. The result is shown in Fig. 4B. It can be observed that the DSINMF achieves the best performance in term of NMI when the dimension is set as 80. In addition, we also test the effect of different layers in deep matrix factorization. The layer of deep matrix factorization is set from 1 to 4. The results are shown in Table 3. It can be found that the performance is the best when the layer is set as 4, i.e four-tire structure matrix factorization.

**TABLE 3.** The performance of DSINMF in Patel with different layers.

| Operating parameters | NMI | AMI | ARI | accuracy |
| --- | --- | --- | --- | --- |
| parameter = (80, k) | 0.7429348 | 0.7504162 | 0.7587987 | 0.8439535 |
| parameter = (80, 70, k) | 0.7633621 | 0.7703931 | 0.7799255 | 0.8613954 |
| parameter = (80,70,60, k) | 0.7747828 | 0.772081333 | 0.8018376 | 0.9269303 |

### 3.3 Application of DSINMF in identifying differentially expressed genes

The cluster results can improve the analysis of downstream scRNA-seq data in the biological field. In here, we aimed to detect significantly differentially expressed genes (DEGs) according to the clustering results. Specifically, the Kruskal Wallis test [29] is used for gene expression profiles with inferred tags. Kruskal Wallis test is a non-parametric method which is usually used to test whether two or more groups come from the same distribution. Then, the DEGs are detected according to the p-value. We focus on the dataset Haber described in Table 1. Haber dataset is a group of 1522 cells from mouse epithelial cells using full length scRNA-seq, consisting of 202 enterocyte cells, 96 enterocyte progenitor early cells, 41 enterocyte progenitor late cells, 637 stem cells, 123 goblet cells, 77 Paneth cells, 102 Tuft cells, 96 enteroendocrine cells and 201 transit amplifying cells (TA). We calculate the p-value of each gene according to the clustering results of DSINMF. If the p-value of a gene meets the significant p-value (p-value <= 0.01), the expression of the gene in one cluster randomly dominates at least another cluster. According to the significant p-value, we selected the top eight genes, which include P2rx4(P=1.41e-121), Fxyd3(P=1.41e-121), Tm4sf4(P=1.41e-121), Selm(P=1.41e-121), Atp2a3(P=1.41e-121), Agr2(P=1.41e-121), Defa24(P=1.41e-121) and Klk1(P=0.0093764).

Fig. 5 shows the changes in the expression levels of top-8 genes on each cell in the Haber dataset. In the figure, it can be clearly seen that the expression value distribution of these genes on various types of cells has obvious differences. For example, the expression of these genes on Paneth cells and Tuft cells almost showed the opposite trend, the genes P2rx4, Tm4sf4, Selm and Atp2a3 show higher expression on Paneth cells, while the genes Agr2, Defa24 and Klk1 show a lower expression level, and the expression trend of these genes on Tuft cells is the opposite. Fig. 6 shows the average expression level of each type of cell on these genes and the sparseness of the gene expression value. It has been proved that the Klk1 plays a key role in the differentiation of goblet and Paneth cells by regulating the biological active of kallikrein [30], [31]. It has been discovered that the Defa24 are responsible for the gene encoding of antimicrobial peptides in mouse Paneth cells that are involved in controlling the intestinal microbiota and immune homeostasis [32]. It has been found that the AGR2 is essential for the production of the intestinal mucin MUC2 in enteroendocrine epithelial cells [33]. It has been proved that the Fxyd3 can reduce glucose-induced insulin secretion by acting downstream of plasma membrane depolarization and Ca2+ influx [34]. Relevant studies have shown that the Tm4sf4 acts as regulation factor in islet progenitor cell during endocrine differentiation [35]. It has been discovered that the Atp2a3 can produces at least seven alternatively spliced SERCA3 subtypes, and the SERCA3 is related to the specific signal regulation of various types of cells in Tuft cells [36].

**Fig. 5.**
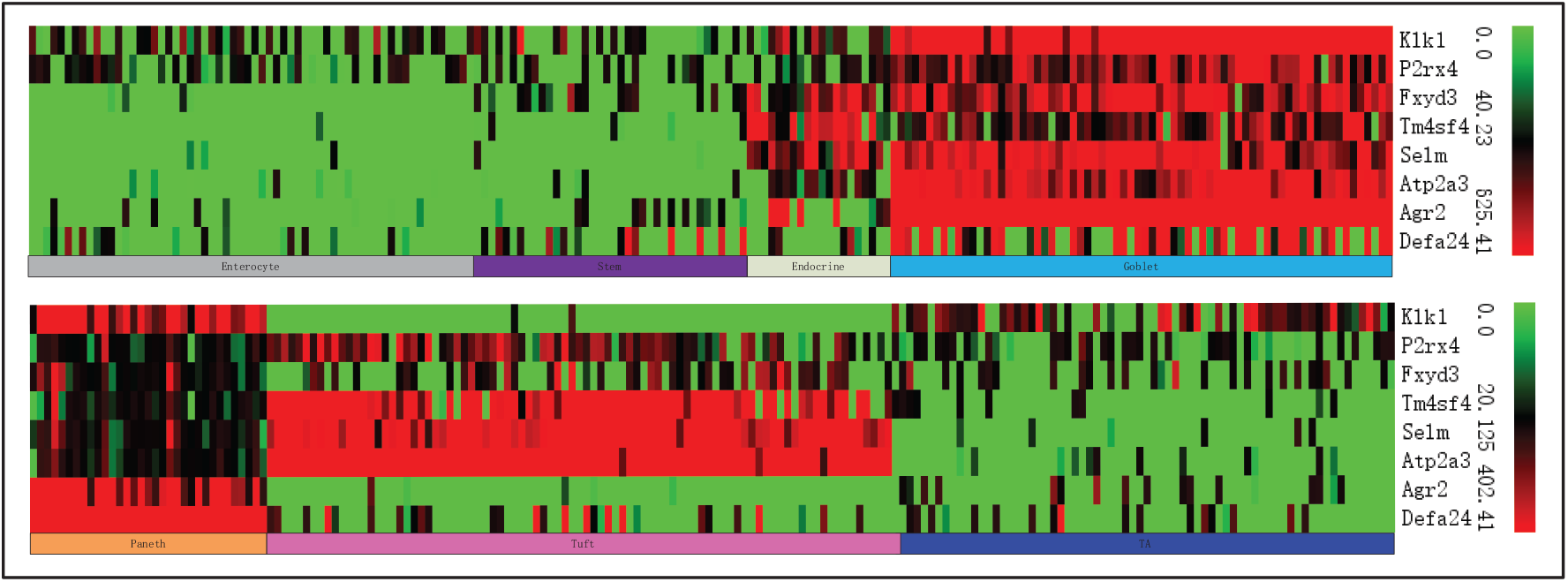
The Heat map of cells in the top-8 gene in the Haber. The Haber dataset has a total of 1,522 cell gene expression data, divided into seven categories.

**Fig. 6.**
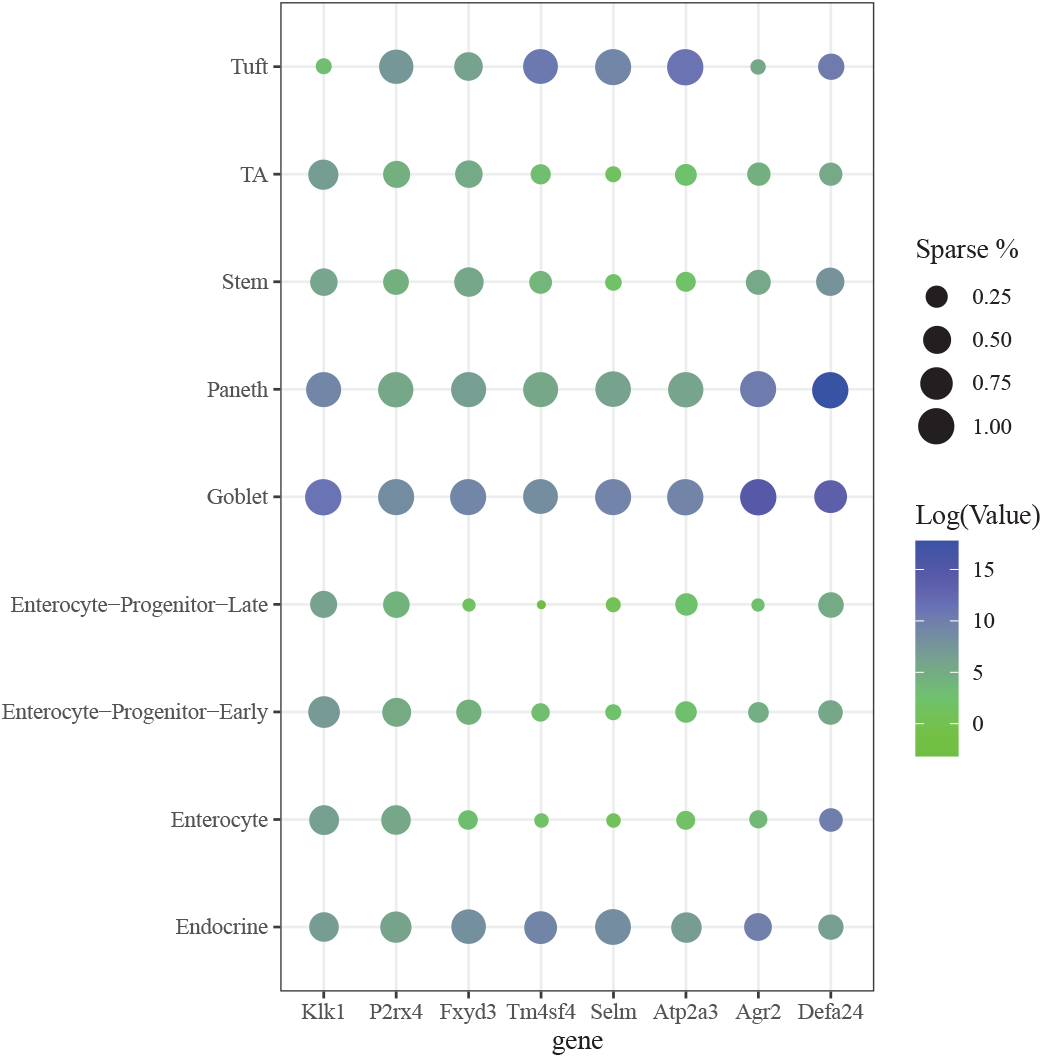
The top 8 gene marker in Haber. The x-axis is the gene name while y-axis is the cell types. The color denotes the expression level of genes and the size of the circles denotes the sparsity of the genes expressing in cells.

## Conclusion

In this paper, we propose a clustering algorithm to cluster scRNA-seq data based on the deep matrix factorization and bi-stochastic graph regularization. In addition, the matrix imputation is utilized to reduce the impact of dropouts. Furthermore, the parameters are learned with adaptive manner. The experimental result show that our method outperformances other state-of-the-art methods. The code is available at http://gxu.biobdlab.cn/code/DSINMF-master.rar and https://github.com/lanbiolab/DSINMF.

## Acknowledgements

This work was partially supported by the National Natural Science Foundation of China (Nos. 62072124, 61963004 and 61972185), the Natural Science Foundation of Guangxi (Nos. 2021GXNSFAA075041 and 2018GXNSFBA281193), the Science and Technology Base and Talent Special Project of Guangxi (No. AD20159044), the Hunan Provincial Science and Technology Program (No. 2018WK4001).

